# Characterisation of Genetically-Derived Patient Clusters Highlights Genetic Heterogeneity of Inflammatory Bowel Disease

**DOI:** 10.64898/2026.10.05.754100

**Authors:** Fernando Vázquez López, Alex Z. Kadhim, Zachary Green, Iman Nazari, Guo Cheng, Louis J Cohen, Judy M Cho, Robert M. Beattie, Sarah Ennis, James J. Ashton

## Abstract

**Background and aims:** Inflammatory Bowel Disease (IBD) is a complex immune-mediated condition with a strong genetic contribution. Its genetics are highly complex, with inflammation, autoimmunity, and mucosal immunity pathways involved in its pathogenicity. GenePy is a gene-level disease burden metric. The integration of clinical and genetic data and the application of machine learning can aid patient stratification for personalised care.

**Methods:** Lists of consensus IBD pathway genes were curated. GenePy scores were computed from whole exome sequencing data. A K-Means algorithm was optimised using clustering quality and stability metrics on the GenePy scores of genes in each IBD pathway, and the results replicated in three independent IBD cohorts (SPARC IBD, MSCCR, and Southampton IBD). Chi-square tests assessed the association between cluster assignment and phenotypic features, and were multiple test corrected via Benjamini-Hochberg FDR.

**Results:** A strong association was replicated for NOD signalling genes driving IBD subtype, Crohn’s disease behaviour, and IBD surgery across all cohorts, as well as granuloma in Southampton IBD. This was mainly driven by *NOD2*. Similar genetic clustering patterns were observed across all cohorts, with additional phenotypic enrichment in clusters driven by *NLRP1* (surgery in MSCCR IBD), and nominally significant enrichment for disease behaviour linked to *BIRC2* (B2 in Southampton IBD and B2B3 in SPARC IBD).

**Conclusions:** This analysis identified a common genetic architecture across NOD signalling genes across IBD cohorts, replicated a signal for *NOD2* as a monogenic disease driver with a strong phenotypic enrichment, and identified other putative genetic drivers of IBD, such as *NLRP1*.

## Introduction

Inflammatory bowel disease (IBD) is a complex inflammatory condition comprised of Crohn’s disease (CD) and ulcerative colitis (UC), which are chronic disorders that affect the gastrointestinal (GI) tract and may cause extra-intestinal manifestations^1,2^. IBD aetiology involves an interplay between genetics, immune dysregulation, and environmental factors such as intestinal dysbiosis. However, the clinical phenotypes and disease complications are extremely heterogenous and go beyond the more reductionist Montreal/Paris classification of disease location and behaviour^3^.

There exists a stronger heritable component in CD than in UC^4^, and genetics plays a more important role in the aetiology of paediatric patients (<18 years at diagnosis) than in adults. Early onset or very early onset cases tend to have a monogenic cause, whereas most individuals diagnosed in adulthood have a more complex polygenic genomic background with a contribution from environmental factors^5,6^. IBD is much more complex at the genetic level than the established UC/CD dichotomy, with cases of monogenic, oligogenic, and polygenic disease, and the identification of both protective and risk variants^7^. This genetic heterogeneity is impinged upon by multiple genes implicated in key molecular pathways. Chronic inflammation, autoimmunity, and mucosal immunity play a major role in the pathogenicity of IBD^7^. The best-established genomic locus in CD is the Nucleotide-binding oligomerization domain-containing protein 2 (*NOD2*)^8,9^. NOD2 plays a key role as a microbial sensor and inflammatory effector in the intestine^10^. Other IBD-relevant biological pathways include antigen presentation and cytokine networks, cellular and oxidative stress, and hypoxia^7,11^. Furthermore, genetic variation in key signalling partners within the same pathway may be causative of different phenotypes and clinical presentations. An example of the aforementioned phenomenon occurs in the NOD signalling pathway, where *NOD2* is an established driver of CD and stricturing^12,13^, and the X-linked inhibitor of apoptosis protein (*XIAP*) gene is involved in immune deficiency and fistulating disease^14,15^. Other genes in this pathway, such as *CHUK, RIPK2, IKBKG*, or *IKBKB* do not have an established IBD-related phenotype. The heterogeneity in the aetiology of specific phenotypes in IBD warrants the identification of stable molecular signatures of disease.

As healthcare provision increasingly focuses on personalised medicine, accurate and clinically actionable patient stratification based on their disease molecular drivers and clinical phenotypes and outcomes becomes more critical. The growing wealth of data available, which includes multi-omics datasets and electronic health records (EHRs) necessitates the development of better methodologies for multi-modal data integration and analysis. However, challenges remain, such as those revolving around data heterogeneity and harmonisation, and lack of suitable tools for processing and analysis^16^. Unsupervised machine learning methods have proved to be very powerful at uncovering associations and structure in high-dimensional data in the context of genetic and clinical data integration. An example of this is work by Chang *et al*., who applied principal components analysis (PCA) and hierarchical agglomerative clustering on the transcriptomes of a UC cohort^17^. Previous work on the Southampton Genetics of IBD cohort used clinical features derived from endoscopic and histological data for PCA and hierarchical clustering^18^.

GenePy is an in-house disease burden metric that summarises germline variant-level pathogenicity into a single score for a given gene or genomic region in an individual. GenePy addresses the sparsity and high dimensionality of genetic data to help analysis and interpretation. Here, we report on a systematic approach for determining consensus lists of genes involved in key IBD biological pathways and implement GenePy and K-Means clustering to characterise the genetic heterogeneity of IBD in three independent patient cohorts. This analysis highlights commonalities in the germline disease burden of IBD genes across independent cohorts and uncovers enrichments for clinical outcomes associated with variation in specific genes.

## Materials and Methods

### IBD Consensus Pathway Analysis

First, a literature review on the current knowledge about pathways/networks implicated in IBD pathogenesis was conducted^7,19,20^. These reviews, and the KEGG (Kyoto Encyclopedia of Genes and Genomes) database Inflammatory Bowel Disease pathway (entry: hsa05321; accessed on 01/08/2024), were used to collate a list of biological processes and signalling networks involved in IBD in humans. The three major pathway databases Reactome (release 89)^21^, WikiPathways^22^, and KEGG (release 111.0)^23^ were searched for entries matching these processes. The online databases were accessed on 01/08/2024. For signalling networks which only had one database source, a literature review was conducted to add a literature source. The goal of this was to have at least two sources per network to curate consensus sets of genes.

Lists of genes and proteins in the chosen pathways from each resource were downloaded: proteins from Reactome, genes and proteins from WikiPathways, and genes from KEGG. Data was downloaded from the databases’ websites, except for KEGG, in which case the REST API was used in a Linux environment. The Ensembl BioMart tool (Ensembl release 112)^24^ was then used to obtain the ”Gene stable ID”, ”Gene description”, ”Gene name”, ”Gene type” and ”UniProtKB Gene Name Symbol” for each gene set. Gene sets were curated to remove duplicates, and a consensus list of genes for each pathway was determined. Where three sources were available, a list of genes in any two sources, and genes in all three was obtained to build sets of high-confidence genes within implicated pathways. Where only two sources were available, gene sets were constructed using either the union or intersection of these.

### Clinical data collection and processing

Within the Southampton Genetics of IBD study, the clinical data of participants, including family history of IBD and history of autoimmune disease, were recorded. Their diagnosis was established in line with the Porto criteria for paediatric cases^25^, and the BSG/ECCO criteria for adults^26,27^. The clinical data for the SPARC IBD and MSCCR cohorts were retrieved from patient electronic health records (EHRs). A large language model (LLM) tool was used to extract phenotypic and clinical information from unstructured clinical records in the Southampton IBD cohort (see Supplementary Methods for details)^28,29^. IBD surgeries which indicated an involvement of the perianal region were used as a proxy for perianal disease.

For the SPARC IBD cohort, patients’ IBD subtype, gender, age at diagnosis, number of IBD surgeries, dates of IBD surgeries, and CD complications were retrieved. All UC and IBDU patients were assigned a B1 label for complications, while CD patients without a status were left as unknown. To harmonise the different sources for IBD surgery, patients with a date of surgery and/or a record of the number of IBD surgeries they had undergone were assigned a ”surgery” status. Patients with no date of surgery and a number of surgeries of 0 were assigned a ”no surgery” status, and any others were assigned an ”unknown” status. For the MSCCR cohort, patients’ IBD subtype, gender, age at diagnosis, number of IBD surgeries, and CD complications were retrieved. The number of surgeries variable was binarised (0: no surgeries; 1: one or more surgeries). The same approach was followed for complications.

### Whole Exome Sequencing and Bioinformatic Analysis

Blood samples were taken from participants in the Southampton Genetics of IBD study and collected in EDTA. Genomic DNA was extracted from these using the salting out method, and Whole Exome Sequencing (WES) was carried out by Novogene, using the Agilent SureSelect human exome capture kit, versions 5 and 6. Illumina paired-end sequencing was used to generate FASTQ files, which were transferred to the Iridis 6 High Performance Computing (HPC) cluster. Details of the procedure for participant recruitment and DNA sequencing can be found in Andreoletti *et al*.^30^. WES data for the SPARC IBD and MSCCR cohorts were available in CRAM format and were extracted to FASTQ for subsequent processing using the same bioinformatic pipeline. Both cohorts used the Twist Custom capture kit. Genetic duplicates were detected and subsequently removed from downstream analysis (four samples in SPARC IBD and 19 in MSCCR).

The exome bioinformatic pipeline used is based on the Genome Analysis Toolkit (GATK) best practices^31^. Where necessary, the raw FASTQ files are merged to produce one forward- and one reverse-read file. Read alignment is performed using the human reference genome (GRCh38 assembly with Human Leukocyte Antigen (HLA) regions) and Burrow-Wheeler Aligned - Maximal Exact Match (BWA-MEM) (bwa version 0.7.18). After alignment, duplicate reads are marked (Picard MarkDuplicates), and base quality recalibration is conducted (GATK BaseRecalibrator and ApplyBQSR) (jdk version 24; GATK version 4.6.2.0; Picard version 3.4.0). The recalibrated BAM files for each sample then undergo variant calling to generate VCF files using GATK HaplotypeCaller, and joint variant calling using GATK GenomicsDBImport and GenotypeGVCF. The joint-call VCF then undergoes further variant recalibration using GATK VariantRecalibrator and ApplyVQSR. Two separate joint-genotyping pipelines were conducted, one for all samples sequenced with an Agilent kit, and the other for all samples sequenced with a Twist kit. The rationale for this was to ameliorate bias derived from the use of different capture kits. The joint-call for SPARC IBD, MSCCR, and the Southampton samples sequenced using Twist Comprehensive was restricted to the union of the genomic targets of ±25 bp Twist Comprehensive with ±25 bp Twist Custom. The joint-call for the Southampton IBD Agilent samples was restricted to the union of ±100 bp Agilent SureSelect V5 and ±100 bp V6. Global ancestry inference was performed using gnomAD pre-computed principal components analysis (PCA) loadings^32^ (see Supplementary Methods for details).

### GenePy Application and Processing

GenePy is a scoring system for pathogenicity or disease burden^33^. This algorithm incorporates variant population allele frequency, zygosity, and a user-defined deleteriousness metric (such as the Combined Annotation Dependent Depletion (CADD) score)^34^. It reduces the dimensionality of variant-level deleteriousness scores to a unique score for any predefined genomic region, and each individual in a cohort. This tool computes a weighted sum of the CADD scores of each locus in an individual, and incorporates population allele frequency from gnomAD 4.1^35^, and variant zygosity. The input for GenePy is a recalibrated joint-call VCF. At the annotation and filtration steps, CADD 1.6 scores are calculated for each variant, and the Ensembl Variant Effect Predictor (VEP – release 114) is used for annotation^36^. Variants are filtered based on variant quality, depth, genotype quality, and missingness. These filtration criteria were derived from analysis by Carson *et al*.^37^. After filtration, variants within exonic Consensus Coding Sequence (CCDS) regions with a padding of ±25 bp and the padded targets of the capture kits included in the joint-call were selected, and gene-annotated with GENCODE V49. The latest release of CCDS regions was downloaded from the UCSC Genome Browser (GRCh38.p14 genome assembly) (release 24 - 27/10/2022). To reduce noise introduced by accumulation of many non-pathogenic variants within a gene, a GenePy matrix for each cohort was computed based on variants with a Phred-scaled CADD score >= 15 (CADD15). This reflects the threshold for likely deleterious variation as defined by CADD. The GenePy pipeline is available from https://github.com/UoS-HGIG/GenePy-2/tree/V2.1/Nextflow_Genepy. Invariant features were removed and the remaining scores min-max scaled for each gene using scikit-learn MinMaxScaler (version 1.7.1; Python version 3.13.5).

### Machine Learning

A K-Means clustering algorithm was implemented on the min-max scaled CADD15 GenePy scores of the selected genes for each IBD cohort, using default hyperparameters. Random Forest Classifiers with random state 1 were implemented to determine the feature importances of each cluster in a one-vs-all manner. To determine the optimal number of clusters (*k*), the algorithm was implemented across 100 iterations for a range of *k* values 2-50. This range was selected to avoid a very large number of clusters, which would result in many patients being assigned to their individual cluster, and reduce the power of statistical testing downstream. For each *k* and each iteration, the Silhouette score and Davies-Bouldin Index (DBI) were computed as measures of internal cluster validity, and the mean pairwise Adjusted Rand Index (ARI) was calculated to quantify clustering stability. A TOPSIS (Technique for Order Preferences by Similarity to an Ideal Solution) based approach was implemented to identify the compromise solution that best satisfies these three optimisation problems (optimising all three metrics). A single per-k per-iteration TOPSIS score was computed, and its median and median absolute deviation across iterations were used to perform local maxima detection. Upon selection of a single optimal k for a set of features in a patient cohort, K-Means was re-implemented with a random state of 42. All machine learning models, and clustering metrics were implemented using the scikit-learn package (see Supplementary Methods for details of the TOPSIS methodology).

### Statistical Analysis

K-Means clusters were annotated with the IBD subtype, CD behaviour, and IBD surgery status for all three cohorts, as well as granuloma, perianal disease, and non-perianal IBD surgeries in the Southampton IBD cohort. All clinical variables were then one-hot encoded for statistical analysis. A chi-square test (χ^2^) of independence was implemented to assess for an association between cluster assignment and each clinical variable. To meet χ^2^ assumptions, any cluster with a count <5 for any class was merged into a “miscellaneous” cluster. Due to the small sample size of patients with an IBDU diagnosis across all cohorts, the count of IBDU patients was dropped from the contingency matrix. The number of patients with a missing value was also dropped for statistical analysis. Individual p-values were calculated from the χ^2^ contributions. Benjamini-Hochberg False Discovery Rate (FDR) was implemented for multiple test correction. Then, standardised residuals were calculated to infer direction. Associations were considered significant at p <0.05. All statistical analysis was conducted in Python, using the scipy (version 1.16.2) and statsmodels (version 0.14.5) packages.

## Results

### Consensus IBD Pathways

Eleven signalling networks were selected due to their involvement in the pathogenicity of IBD (Table 1). It should be noted that these pathways are not completely independent, but rather, their signalling networks overlap reflecting the complex interactions between different parts of the innate and adaptive immune system. For some of the pathways, only a very small number of genes were present in all sources (such as IL-10 signalling, IL-23 signalling, or JAK-STAT signalling).

**Table 1.**
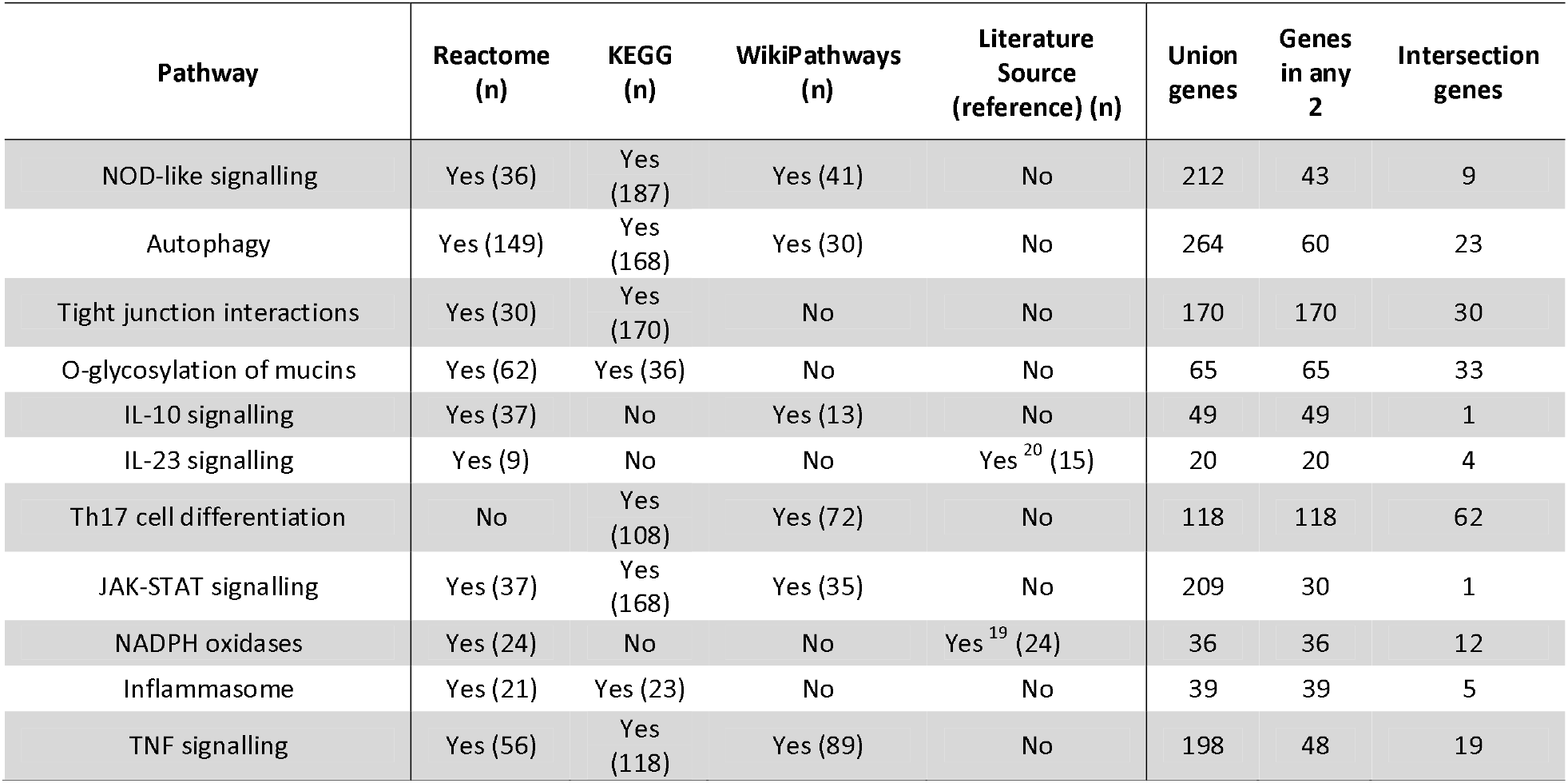
Results from curation of consensus gene sets in IBD pathways. Sources from the literature are provided when applicable. NOD: Nucleotide-binding Oligomerisation Domain; KEGG: Kyoto Encyclopaedia of Genes and Genomes; IL: Interleukin; JAK-STAT: Janus Kinase-Signal Transducer and Activator of Transcription; NADPH: Nicotinamide Adenine Dinucleotide Phosphate; TNF: Tumour Necrosis Factor.

### IBD Cohorts

Three independent IBD cohort were included in the analysis. See Table 2 for demographic and clinical characteristics. Supplementary Table 1 summarises genetically inferred ancestry. The ‘Genetics of Paediatric IBD Study – Southampton’ is an approved portfolio study (Regional ethics committee approval number-REC 09/H0504/125). Patients were recruited through the gastroenterology services at University Hospital Southampton (UHS), UK. SPARC (Study of a Prospective Adult Research Cohort) IBD is a disease-only cohort from the Crohn’s and Colitis Foundation’s IBD Plexus Program in the US^38,39^. The Mount Sinai Crohn’s and Colitis Registry (MSCCR) is a US-based cohort that includes IBD and control patients^39,40^.

**Table 2.** Demographic characteristics of IBD cohorts included in the analysis. In SPARC IBD, 37 patients did not have an age at diagnosis available. In the Southampton IBD cohort, three patients did not have an age at diagnosis available, and seven patients did not have a CD/UC/IBDU diagnosis. SPARC: Study of a Prospective Adult Research Cohort; MSCCR: Mount Sinai Crohn’s and Colitis Registry.

| Cohort | All participants with IBD | CD | UC | IBDU | Males | Females | Median age at diagnosis | Ratio of paediatric-to-adult onset |
| --- | --- | --- | --- | --- | --- | --- | --- | --- |
| Southampton IBD | 1,724 | 1,044 | 609 | 71 | 883 | 841 | 18.59 | 827:900 (0.92) |
| SPARC IBD | 2,856 | 1,900 | 906 | 50 | 1,286 | 1,570 | 25 | 617:2,202 (0.28) |
| MSCCR IBD | 1,038 | 586 | 441 | 11 | 564 | 474 | 25 | 207:831 (0.25) |

### Clinical data availability

Some IBD patients with WES data did not have complete clinical records. Supplementary Table 2 provides a summary of the availability of clinical data in each cohort.

### K-Means Clustering and χ^2^ Analysis Consistently Confirm a Signal for Variation in NOD Signalling Genes Driving CD, Stricturing, and Surgery

The GenePy scores of consensus genes in the aforementioned key IBD signalling pathways were used to optimise K-Means clustering models in the Southampton IBD, SPARC IBD, and MSCCR IBD cohorts. For each pathway, only genes present in all available sources were used, and features which were available in all cohorts were included in the analysis. For the IL-10 signalling and JAK-STAT signalling pathways, only one gene was selected following this method, and they were not included in the analysis. To avoid noise from a large number of highly sparse features and poor K-Means performance Th17 cell differentiation genes were also excluded.

The optimal k for each of these pathways and cohorts was selected using TOPSIS-based local maxima detection, K-Means was re-implemented using a set random state, and the resulting clusters were annotated with the patients’ IBD subtype, IBD surgery and CD complications (stricturing and fistulating disease). Further clinical outcome data on perianal disease and granuloma was only available for patients in the Southampton IBD cohort. Supplementary Table 3 shows the number of clusters selected for each gene set and each cohort, and the results from χ^2^ testing.

The GenePy scores of eight NOD signalling genes (*CARD9, CASP1, CASP8, CHUK, IKBKB, NOD1, NOD2*, and *RIPK2*) yielded the most significant associations between cluster assignments and clinical outcomes. GenePy scores were not available for the *IKBKG* gene in all cohorts. For detailed results of k optimisation see Supplementary Table 4. There was a significant association between cluster assignment and IBD subtype (CD and UC), CD behaviour (stricturing and fistulating disease), and IBD surgery in all cohorts. Furthermore, NOD signalling genes drove an association with non-perianal IBD surgery and granuloma in the Southampton IBD cohort (Table 3).

**Table 3.** χ^2^ results for the association between cluster assignment and clinical outcomes in three IBD cohorts. χ2 tests were implemented to assess the association between cluster assignment and IBD subtype, CD behaviour, and IBD surgery for all cohorts. Tests were also performed for perianal disease, non-perianal surgery, and granuloma in the Southampton IBD cohort. A K-Means clustering algorithm was implemented using the min-max scaled CADD15 GenePy scores of eight NOD signalling genes. NOD: Nucleotide-binding Oligomerisation Domain; SPARC: Study of a Prospective Adult Research Cohort; MSCCR: Mount Sinai Crohn’s and Colitis Registry.

| Cohort | Number of samples | K | Max cluster size | $\chi^2$ IBD subtype p-value | $\chi^2$ CD behaviour p-value | $\chi^2$ IBD surgery p-value | $\chi^2$ perianal disease p-value | $\chi^2$ non-perianal IBD surgery p-value | $\chi^2$ granuloma p-value |
| --- | --- | --- | --- | --- | --- | --- | --- | --- | --- |
| SPARC IBD | 2,856 | 22 | 608 | 8.63E-15 | 5.74E-10 | 2.31E-05 | - | - | - |
| MSCCR IBD | 1,038 | 8 | 604 | 2.17E-04 | 1.51E-04 | 4.41E-08 | - | - | - |
| Southampton IBD | 1,724 | 13 | 548 | 6.40E-09 | 6.07E-05 | 1.07E-02 | 6.87E-01 | 2.00E-02 | 3.01E-03 |

These clusters are illustrated as heatmaps, annotated by the GenePy scores of each feature, and the clinical outcomes (Figure 1). Some similarities between the structure driven by the GenePy scores of these genes in each cohort are apparent. Most patients had a non-zero GenePy score (indicative of them carrying a Phred-scaled CADD ≥15 variant) for *CASP8, NOD1* and *CHUK* across all cohorts. The *CASP8* and *IKBKB* genes both drove one small cluster in each cohort. Variation in the *NOD2* gene drove patients into well-defined and distinct clusters. The significance of cluster-level enrichment or deviation for any clinical outcome was also calculated and multiple test corrected using Benjamini-Hochberg FDR. Analysis of these results identified a strong signal from variants in the *NOD2* gene driving Crohn’s disease, stricturing, fistulating disease and surgery. However, some of these enrichments did not withstand multiple test correction (Table 4). The *NOD2* gene also showed a signal for granuloma and non-perianal IBD surgeries (Table 5).

**Table 4.** Cluster-level χ^2^ results for the association between cluster assignment and clinical outcomes in three IBD cohorts. χ^2^ tests were implemented to assess the association between cluster assignment and IBD subtype, Crohn’s disease behaviour, and IBD surgery. A K-Means clustering algorithm was implemented using the min-max scaled CADD15 GenePy scores of eight NOD signalling genes. Only clusters with a nominally significant p-value for any clinical outcome have been included. “^*^” denotes a nominally significant enrichment or depletion for a clinical outcome. Bolding denotes these p-values withstood Benjamini-Hochberg False Discovery Rate (FDR) multiple test correction. No patients in MSCCR had a CD complication diagnosis of B2B3 (stricturing and fistulating disease). SPARC: Study of a Prospective Adult Research Cohort; MSCCR: Mount Sinai Crohn’s and Colitis Registry.

| Cohort (k) | Cluster | Number of samples | GenePy drivers | UC | CD | B1 | B2 | B3 | B2B3 | No surgery | Surgery |
| --- | --- | --- | --- | --- | --- | --- | --- | --- | --- | --- | --- |
| SPARC IBD (k = 22) | 1 | 183 | *NOD2* | **3.4E-05\*** | **4.2E-03\*** | **1.9E-04\*** | **8.0E-05\*** | 1.9E-02* | **3.0E-03\*** | 1.0E-01 | 5.1E-02 |
|  | 2 | 75 | *RIPK2; NOD2* | 4.3E-02* | 1.6E-01 | 2.8E-01 | 2.0E-01 | 9.2E-01 | 1.7E-01 | 1.5E-01 | 8.8E-02 |
|  | 7 | 467 | *NOD2* | **4.6E-03\*** | 5.0E-02 | 2.7E-01 | 5.6E-02 | 4.6E-01 | 7.0E-01 | 2.9E-01 | 2.1E-01 |
|  | 11 | 255 | *RIPK2; NOD2* | 1.3E-02* | 8.5E-02 | 3.1E-01 | 1.6E-01 | 6.6E-01 | 6.1E-01 | 4.6E-01 | 3.8E-01 |
|  | 12 | 608 | *NOD1; NOD2; RIPK2; CHUK* | 4.2E-02* | 1.6E-01 | 7.3E-02 | **2.2E-03\*** | 4.7E-01 | 9.6E-01 | 3.9E-01 | 3.1E-01 |
|  | 14 | 68 | *NOD2* | 1.0E-01 | 1.6E-01 | 2.6E-01 | 7.9E-01 | 1.8E-02* | 2.6E-01 | 2.1E-01 | 1.3E-01 |
|  | 16 | 359 | *NOD2* | **4.6E-03\*** | 5.0E-02 | 1.4E-01 | 7.0E-01 | 1.9E-02* | 2.1E-01 | 9.3E-02 | 4.6E-02* |
| MSCCR (k = 9) | 0 | 604 | *NOD2; RIPK2* | 1.4E-01 | 9.1E-02 | 2.5E-01 | 6.0E-02 | 3.7E-01 | NA | 1.6E-01 | 1.6E-02* |
|  | 1 | 119 | *NOD2* | 3.7E-03* | 1.2E-02* | 3.3E-02* | **4.7E-04\*** | 1.0E-01 | NA | 9.8E-03* | **1.2E-05\*** |
| Southampton IBD (k = 13) | 0 | 288 | *NOD2* | 1.7E-02* | 6.9E-02 | 2.4E-01 | 6.3E-01 | 2.7E-01 | 4.0E-02* | 2.6E-01 | 3.5E-02* |
|  | 2 | 548 | *NOD1; NOD2; RIPK2* | 2.0E-01 | 3.3E-01 | 5.1E-01 | 3.1E-01 | 2.4E-01 | 3.1E-02* | 7.3E-01 | 5.2E-01 |
|  | 5 | 149 | *NOD2* | **4.0E-06\*** | **4.3E-04\*** | 3.5E-02* | **1.7E-04\*** | 7.8E-01 | 4.5E-02* | 2.3E-01 | 2.6E-02* |
|  | 7 | 412 | *NOD1; NOD2; RIPK2* | 5.0E-02* | 1.3E-01 | 3.2E-01 | 9.2E-02 | 2.4E-01 | 6.3E-01 | 4.1E-01 | 1.2E-01 |

**Table 5.** Cluster-level χ^2^ results for the association between cluster assignment and additional clinical outcomes in the Southampton IBD cohort. χ2 tests were implemented to assess the association between cluster assignment and perianal disease, non-perianal surgery, and granuloma. A K-Means clustering algorithm was implemented using the min-max scaled CADD15 GenePy scores of consensus IBD signalling genes. Only clusters with a nominally significant p-value for any clinical outcome have been included. “^*^” denotes a nominally significant enrichment or depletion for a clinical outcome.

| Cohort (k) | Cluster | Number of samples | GenePy drivers | No perianal disease | Perianal disease | No non-perianal surgery | Non-perianal surgery | No granuloma | Granuloma |
| --- | --- | --- | --- | --- | --- | --- | --- | --- | --- |
| Southampton IBD (k = 13) | 0 | 288 | *NOD2* | 9.9E-01 | 9.7E-01 | 2.9E-01 | 3.5E-02 | 5.3E-01 | 3.9E-01 |
|  | 5 | 149 | *NOD2* | 8.1E-01 | 2.9E-01 | 3.6E-01 | 6.8E-02 | 1.2E-01 | 3.2E-02 |

**Figure 1.**
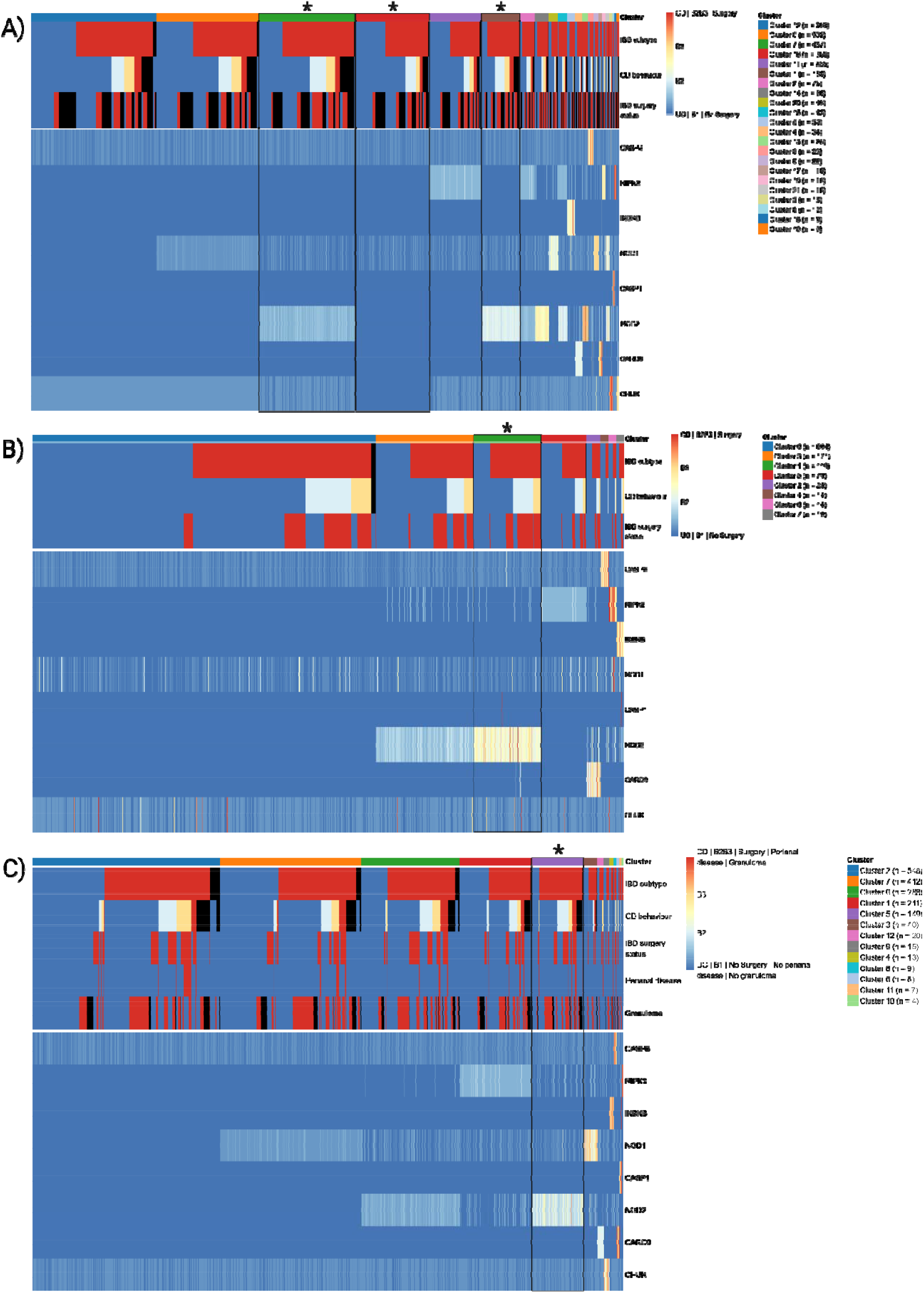
Heatmaps of K-Means clusters on the min-max scaled CADD15 GenePy scores of 8 NOD signalling genes in three IBD cohorts. A) SPARC IBD (n = 2,856). B) MSCCR IBD (n = 1,038). C) Southampton IBD (n = 1,724). Clusters are sorted by size and have been annotated with the patients’ clinical outcomes (top panel) and the GenePy scores of the features used (bottom panel). The colour black denotes an IBDU diagnosis for the IBD subtype, and missing data for Crohn’s disease behaviour, IBD surgery, perianal disease and granuloma. “^*^” denote clusters with a significant enrichment and/or depletion for a clinical outcome after multiple test correction.

### K-Means Clustering and χ^2^ Uncover the *NLRP1* Gene as a Putative Driver of Surgery in IBD

As shown by Supplementary Table 3, the associations with clinical outcomes which reached statistically significance in the other signalling pathways were markedly less strong than those driven by NOD signalling genes. Within the MSCCR IBD cohort, there was a significant association between cluster assignment and IBD surgery status (p = 0.011). Even though the enrichment or depletion for clinical outcomes of most clusters driven by non-NOD signalling genes could not withstand multiple test correction, within this clustering, cluster 10 had a significant enrichment for surgery which withstood multiple test correction and was driven by genetic variation in the *NLRP1* gene (Table 6).

**Table 6.**
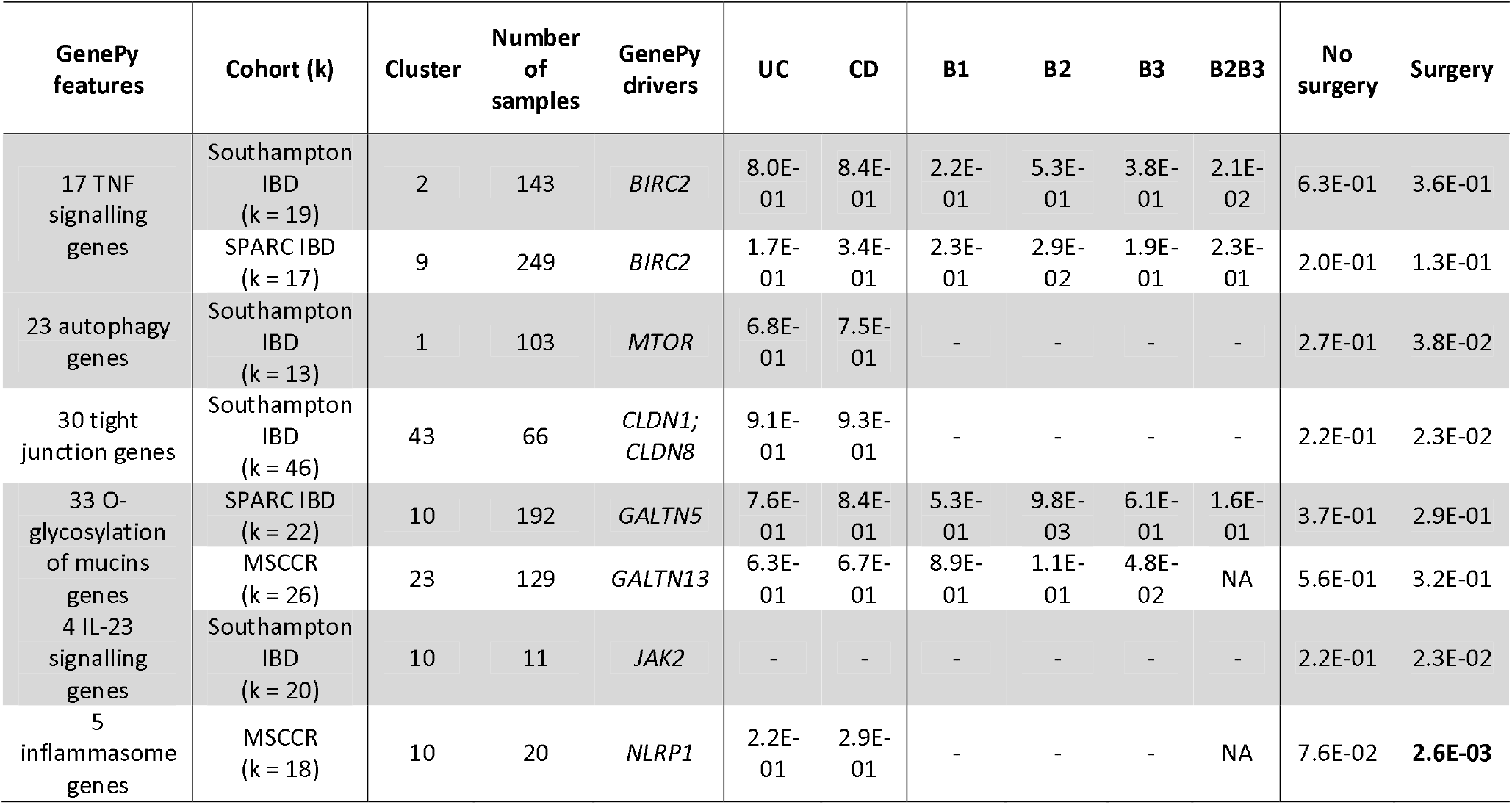
Cluster-level χ^2^ results for the association between cluster assignment and clinical outcomes in three IBD cohorts. χ2 tests were implemented to assess the association between cluster assignment and IBD subtype, Crohn’s disease behaviour, and IBD surgery. A K-Means clustering algorithm was implemented using the min-max scaled CADD15 GenePy scores of consensus IBD signalling genes. Only clusters with a nominally significant p-value for any clinical outcome have been included. “^*^” denotes a nominally significant enrichment or depletion for a clinical outcome. Bolding denotes these p-values withstood Benjamini-Hochberg False Discovery Rate (FDR) multiple test correction. No patients in MSCCR had a CD complication diagnosis of B2B3 (stricturing and fistulating disease). In cases where χ2 assumptions could not be met (all counts in the contingency matrix must be >5), the test was not performed, and this is denoted by “-“. TNF: Tumour Necrosis Factor; IL: Interleukin; SPARC: Study of a Prospective Adult Research Cohort; MSCCR: Mount Sinai Crohn’s and Colitis Registry.

The 20 patients in this cluster were investigated further. 13 of them had the same genetic variant behind their GenePy score (s73973843; NC 000017.11:g.5558895T>A). According to gnomAD 4.1, this variant has different ancestry-specific allele frequencies across genomes and exomes; the allele frequency for non-Finnish Europeans was 0.0002, but 0.092 for African/African American individuals. Eight of the 13 individuals carrying this variant had a predicted African/African American genetic ancestry, and the remaining five were admixed.

## Discussion

In this work, we report on the application of a K-Means algorithm and a TOPSIS-based local maxima detection methodology to optimise clustering of GenePy data. This approach was replicated in three independent IBD cohorts to investigate the commonalities and differences in the genetic architecture driven by variants in consensus IBD signalling genes. Clinical outcome data were integrated to identify significant associations with cluster assignment. We confirmed a strong signal from genetic variation in NOD signalling genes (especially *NOD2*) driving an enrichment for CD, stricturing disease, surgery, and granuloma in IBD patients. We also identified other rarer putative genetic drivers of specific disease outcomes, in particular the *NLRP1* gene.

The TOPSIS-based local maxima detection approach provided a systematic way of selecting an optimal k in K-Means clustering. By integrating the clustering quality metrics silhouette score and DBI with the ARI for clustering stability, this addresses discrepancies between metrics and provides a single score per value of k. Clusterings driven by the GenePy scores of 8 NOD signalling genes had by some margin the most significant χ2 associations with clinical outcomes (Table 3 and Supplementary Table 3). This recapitulates a well-known signal for variants in the *NOD2* gene driving monogenic Crohn’s disease, and higher rates of stricturing and surgery^12,41^. The link between *NOD2* variants and fistulating disease and granuloma is less well-established. Work by Schnitzler *et al*. identified a single nucleotide polymorphism (SNP) in *NOD2* as a predictor of perianal fistulas in CD patients^42^. Despite some signal from the *RIPK2* and *NOD1* genes, the association seems to be mainly driven by *NOD2*. While the similarities between the clusterings driven by these genes across cohorts are suggestive of putative common disease drivers, the small size of many of these clusters (such as those driven by the GenePy scores of *CASP8, RIPK2, IKBKB* and *CARD9*) limits statistical power.

The TOPSIS method appeared to perform worse on a smaller number of features. It could not identify a local maximum using inflammasome genes for any cohort, nor using IL-23 signalling genes in SPARC IBD. In clusterings driven by either of these feature sets, most patients were assigned to the same cluster (Supplementary Table 3). This may be due to most patients in these cohorts not having any CADD15 variants in these genes. Significant χ2 associations with clinical outcomes were markedly less strong than those from NOD signalling genes (Supplementary Table 3). An explanation for this may be that most genes involved in monogenic, oligogenic or polygenic IBD are relatively rare drivers; whereas *NOD2* is one of the most common genetic causes of CD. This may also explain why many individual clusters with a nominally significant enrichment or depletion for a clinical outcome could not withstand multiple test correction (Table 6). One hypothesis for the *NLRP1*-surgery association in the MSCCR cohort is an ancestry-specific risk locus or gene-disease association among individuals with an African or African American genetic background.

There are some limitations to the analysis performed in this chapter. First, clinical data missingness among IBD patients limits statistical power and may overestimate or underestimate associations. Due to the lack of a true control cohort free of autoimmune disease with available genetic and clinical data, clustering was performed on patients with a confirmed IBD diagnosis only. Therefore, the genetic structure and associations with clinical outcomes among patients could not be compared against controls. A trade-off emerged by which a higher number of clusters correlated with better clustering quality and stability (as reported by TOPSIS scores) and statistical power. A higher k may also uncover genes driving small well-defined clusters of patients. However, a higher number of clusters will increase the size of the contingency matrix in χ2, thus increasing the penalty from multiple test correction. Finally, patients in these cohorts were joint-genotyped in two different groups based on WES capture kits, and despite GenePy score harmonisation some batch effects may remain downstream.

An alternative approach to cluster patients based on their GenePy scores would be to aggregate gene-level GenePy scores across consensus genes in a signalling network to compute aggregated pathway GenePy scores (i.e. NOD signalling, JAK-STAT signalling, etc.). However, this may hinder our ability to separate patients and would negatively impact model interpretability. We also chose not to test the union of gene sets in different IBD signalling pathways to prevent the negative impact on K-Means performance from a very high number of GenePy features, which tend to be low variance and highly sparse. Furthermore, this would have again limited interpretability, and statistical power.

## Acknowledgments

The authors acknowledge the use of the Iridis 6 High Performance Computing cluster for bioinformatic analysis, the Southampton Emerging Technologies and Therapies (SETT) data and AI team, and the Clinical Informatics Research Unit (CIRU). The results published here are partly based on data from the Study of a Prospective Adult Research Cohort with IBD (SPARC IBD). SPARC IBD is a component of the Crohn’s & Colitis Foundation’s IBD Plexus data exchange platform. Above all, we would like to thank the patients and their families.

## Data Availability Statement

The SPARC IBD data are available upon approved application to Crohn’s & Colitis Foundation IBD Plexus (https://www.crohnscolitisfoundation.org/ibd-plexus). Data from the Southampton Genetics of Inflammatory Bowel Disease study can be shared via appropriate data sharing arrangements.

## Notes

### Competing Interest Statement

JJA is a scientific advisory board member for Orchard Therapeutics, he has contributed to a personalised IBD think tank (Takeda funded) and participated in a IBD transition Delphi consensus process (Pfizer funded). The other authors declare no conflicts of interest in relation to the submitted work.

