## Supplementary Methods for "Characterisation of Genetically-Derived Patient Clusters Highlights Genetic Heterogeneity of Inflammatory Bowel Disease"

Alex Z. Kadhim^1,2^ PhD

Zachary Green^1,3^

Iman Nazari^1^ PhD

Guo Cheng^1,2^ PhD

Louis J Cohen^4^ MD

Judy M Cho^5^ MD

Robert M. Beattie^3^ FRCPCH

Sarah Ennis^1,2^ PhD

James J. Ashton^1,3^ PhD

^1^Human Genetics and Genomic Medicine, University of Southampton, Southampton, UK

^2^National Institute for Health Research (NIHR) Southampton Biomedical Research Centre, Southampton, UK

^3^Department of Paediatric Gastroenterology, University Hospital Southampton, UK

^4^Division of Gastroenterology, Icahn School of Medicine at Mount Sinai, New York, US

^5^Department of Genetics & Genomic Sciences, Charles Bronfman Institute for Personalized Medicine, Icahn School of Medicine at Mount Sinai, New York, US

**Supplementary Methods:**

**Clinical data collection and processing:**

For the Southampton IBD cohort, a novel, in-house Large Language Model (LLM), based on the OpenAI open-weight gpt-oss-120b model, was used to retrieve phenotypic and clinical information from unstructured clinical records^1,2^. The LLM extracted statements from histology and radiology reports’ clinical history, pathology, diagnosis, and treatment sections, as previously described^1,2^. The structured output (finding and region pairs) from this tool was restricted to anatomical regions in the GI tract. Statements from the reports’ pathology sections were queried for instances of granuloma (histology only) and stricturing or fistulating disease (radiology only). This was due to imaging not being suitable to detect the presence of granulomas. The established nomenclature for CD complications is: B1 (inflammation without presence of stricture or fistulae), B2 (inflammation with stricture), B3 (inflammation with penetrating disease), and B2B3 (inflammation with stricture and penetrating disease).

**Prediction of gnomAD Genetic Ancestries:**

gnomAD ancestries were inferred using pre-computed principal components analysis (PCA) loadings from v4.0. Joint-call VCFs were normalised to split multi-allelic sites using bcftools norm and restricted to sites common to both. These processed VCFs were then imported as hail tables, temporarily converted to VariantDataset (VDS) format to filter for sites in the loadings table, densified back into hail tables, and further filtered by GQ >=20, DP >= 10, and het AB >=0.2 and <=0.8 (Python 3.11.15; Hail 0.2.137; gnomad 0.8.2). Genotypes were projected onto the loadings using Hail, and a onnx random forest classifier model was implemented to determine ancestry probabilities, using the minimum required 20 principal components and a minimum probability for ancestry assignment of 0.8 (80%) (onnx 1.20.0). Probabilities were computed for nine ancestries: African/African American (AFR), Admixed American (AMR), East Asian (EAS), Middle Eastern (MID), South Asian (SAS), Amish (AMI), Ashkenazi Jewish (ASJ), Finnish (FIN), and non-Finnish European (NFE). Individuals with no probability above or equal to the minimum probability were classed as “other”. The PCA loadings and random forest model are publicly available for download at <https://gnomad.broadinstitute.org/data#v4-genetic-ancestry-group-classification>.

**TOPSIS-based Optimisation of the Number of Clusters in K-Means:**

The TOPSIS (Technique for Order Preferences by Similarity to an Ideal Solution) method was first proposed by Hwang and Yoon (1981) to determine the best alternative based on the concepts of the compromise solution^3^. A compromise solution is the solution with the shortest Euclidean distance from the ideal solution and farthest Euclidean distance from the negative ideal solution^4^. This method was used to address the Multi-Criteria Decision Analysis (MCDA) problem of determining the optimal k from Silhouette, DBI, and mean pairwise ARI values.

Given a set of alternatives $K={\{K}_{k}\left| k=2,\ldots,n \right\};n=50$, a set of criteria (clustering quality and stability metrics) $C=\{C_{i}|j=SIL,DBI,PARI\}$ (where SIL denotes the Silhouette score, DBI the Davies-Bouldin Index, and PARI the pairwise Adjusted Rand Index), and a set of weights $w=\left\{ wj \right\}=[0.33, 0.33, 0.33]$, a decision matrix is built and ratings are normalised. Normalisation compresses all three metrics to the same range [0, 1]. These normalised ratings were weighted, and then the ideal and negative ideal points for TOPSIS were defined:

$$PIS=v_{j}^{+}=\max_{k} v_{k,j}$$

$$NIS=v_{j}^{-}=\min_{k} v_{k,j}$$

From these Euclidean distances, closeness coefficients to the ideal point were computed for each alternative:

$${C_{k}^{+} = D}_{k}^{-}/\left( D_{k}^{+}+D_{k}^{-} \right),k=1,...,n$$

Closeness coefficients (TOPSIS scores) were calculated for each iteration and each value of k, and their median, median absolute deviation, and 95% confidence intervals (CI) were computed for each k across all iterations.

These TOPSIS scores were then subjected to local maxima detection to optimise the number of clusters in K-Means. The use of local maxima detection rather than standard ranking of scores is justified by the non- monotonic nature of a TOPSIS score distribution computed from silhouette, DBI, and pairwise ARI values (while both the silhouette score and the ARI tend to increase with k; this is not the case for the DBI). The following algorithm was used:

1. Select local maxima within a local window n = 2. This returns values of k which have the highest median TOPSIS score within ± 2 k.
2. Compute prominence of selected k values (height of median TOPSIS above average of median TOPSIS scores within the local window).
3. Apply prominence cut-off of 0.01 (TOPSIS has a theoretical range of [0, 1] due to pre-computation scaling of clustering metrics).
4. Subtract TOPSIS scores median absolute deviation from median in remaining candidates to penalise variability and select k with the highest score.
5. If no candidate k remains, filter for a local window n = 1 with a prominence cut-off of 0.01.
6. If still no candidate k remains, filter for a local window n = 1 with a prominence cut-off of 0.001.
7. Else, select k with highest median TOPSIS score minus standard deviation. Where local maxima detection fails to find any candidate k values, the algorithm defaults to ranking.

**List of References:**

1.	Kadhim, A. Z. *et al.* Application of generative artificial intelligence to utilize unstructured clinical data for acceleration of inflammatory bowel disease research. *Med* 100895 (2025) doi:10.1016/j.medj.2025.100895.

2.	Kadhim, A. Z. *et al.* Human Phenotype Ontology (HPO) Mapper: Semantic Mapping of Clinical Findings to the Human Phenotype Ontology Using AI-Powered Embeddings and LLM-Based Quality Control. Preprint at https://doi.org/10.64898/2025.12.20.25342726 (2025).

3.	Hwang, C.-L. & Yoon, K. Multiple Attributes Decision Making Methods and Applications. *Multiple Attribute Decision Making* (1981).

4.	Tzeng, G. H. & Huang, J. J. *Multiple Attribute Decision Making: Methods and Applications*. *Multiple Attribute Decision Making: Methods and Applications* (2011).
